# GeneSeqToFamily: the Ensembl Compara GeneTrees pipeline as a Galaxy workflow

**DOI:** 10.1101/096529

**Authors:** Anil S. Thanki, Nicola Soranzo, Wilfried Haerty, Robert P. Davey

## Abstract

**Background:** Gene duplication is a major factor contributing to evolutionary novelty, and the contraction or expansion of gene families has often been associated with morphological, physiological and environmental adaptations. The study of homologous genes helps us to understand the evolution of gene families. It plays a vital role in finding ancestral gene duplication events as well as identifying genes that have diverged from a common ancestor under positive selection. There are various tools available, such as MSOAR, OrthoMCL and HomoloGene, to identify gene families and visualise syntenic information between species, providing an overview of syntenic regions evolution at the family level. Unfortunately, none of them provide information about structural changes within genes, such as the conservation of ancestral exon boundaries amongst multiple genomes. The Ensembl GeneTrees computational pipeline generates gene trees based on coding sequences and provides details about exon conservation, and is used in the Ensembl Compara project to discover gene families.

**Findings:** A certain amount of expertise is required to configure and run the Ensembl Compara GeneTrees pipeline via command line. Therefore, we have converted the command line Ensembl Compara GeneTrees pipeline into a Galaxy workflow, called GeneSeqToFamily, and provided additional functionality. This workflow uses existing tools from the Galaxy ToolShed, as well as providing additional wrappers and tools that are required to run the workflow.

**Conclusions:** GeneSeqToFamily represents the Ensembl Compara pipeline as a set of interconnected Galaxy tools, so they can be run interactively within the Galaxy’s user-friendly workflow environment while still providing the flexibility to tailor the analysis by changing configurations and tools if necessary. Additional tools allow users to subsequently visualise the gene families produced by the workflow, using the Aequatus.js interactive tool, which has been developed as part of the Aequatus software project.

## Introduction

The phylogenetic information inferred from the study of homologous genes helps us to understand the evolution of gene families, which plays a vital role in finding ancestral gene duplication events as well as identifying regions under positive selection within species [1]. In order to investigate these low-level comparisons between gene families, the Ensembl Compara GeneTrees gene orthology and paralogy prediction software suite [2]was developed as a pipeline that uses TreeBest [3] [4] (part of TreeFam [5]) to find internal structural-level synteny for homologous genes. TreeBeST implements multiple independent phylogenetic methods and can merge their results in a consensus tree whilst trying to minimise duplications and deletions relative to a known species tree. This allows TreeBeST to take advantage of the fact that DNA-based methods are often more accurate for closely related parts of trees, while protein-based trees are better at longer distances.

The Ensembl GeneTrees pipeline comprises seven basic steps, starting from a set of protein sequences and performing similarity searching and multiple large-scale alignments to infer homology among them, using various tools: BLAST [6], hcluster_sg [7], T-Coffee [8], and phylogenetic tree construction tools, including TreeBeST. Whilst all these tools are freely available, most are specific to certain computing environments, are only usable via the command line, and require many dependencies to be fulfilled. Therefore, users are not always sufficiently expert in system administration in order to install, run, and debug the various tools at each stage in a chain of processes. To help ease the complexity of running the GeneTrees pipeline, we have employed the Galaxy bioinformatics analysis platform to relieve the burden of managing these system-level challenges.

Galaxy is an open-source framework for running a broad collection of bioinformatics tools via a user-friendly web interface [9]. No client software is required other than a recent web browser, and users are able to run tools singly or aggregated into interconnected pipelines, called *workflows*. Galaxy enables users to not only create, but also share workflows with the community. In this way, it helps users who have little or no bioinformatics expertise to run potentially complex pipelines in order to analyse their own data and interrogate results within a single online platform. Furthermore, pipelines can be published in a scientific paper or in a repository such as myExperiment [10] to encourage transparency and reproducibility.

In addition to analytical tools, Galaxy also contains plugins [11] for data visualisation. Galaxy visualisation plugins may be interactive and can be configured to visualise various data types, for example, bar plots, scatter plots, and phylogenetic trees. It is also possible to develop custom visualisation plugins and easily integrate them into Galaxy. As the output of the GeneSeqToFamily workflow is not conducive to human readability, we also provide a data-to-visualisation plugin based on the Aequatus software [12]. Aequatus.js [13] is a new JavaScript library for the visualisation of homologous genes, which is extracted from the standalone Aequatus tool. It provides a detailed view of gene structure across gene families, including shared exon information within gene families alongside gene tree representations. It also shows details about the type of interrelation event that gave rise to the family, such as speciation, duplication, and gene splits.

## Methods

The GeneSeqToFamily workflow has been developed to run the Ensembl Compara software suite within the Galaxy environment, combining various tools alongside preconfigured parameters obtained from the Ensembl Compara pipeline to produce gene trees. Among the tools used in GeneSeqToFamily (listed in Table 1), some were existing tools in the Galaxy ToolShed [14], such as NCBI BLAST, TranSeq, Tranalign and various format converters. Additional tools that are part of the pipeline were developed at the Earlham Institute (EI) and submitted to the ToolShed, i.e. *BLAST parser, hcluster_sg, hcluster_sg parser, T-Coffee, TreeBeST best* and *Gene Alignment and Family Aggregator*. Finally, we developed helper tools that are not part of the workflow itself, but aid the generation of input data for the workflow and these are also in the ToolShed, i.e. *Get features by Ensembl ID, Get sequences by Ensembl ID, Select longest CDS per gene, ETE species tree generator* and *GeneSeqToFamily preparation*.

**Table 1:**
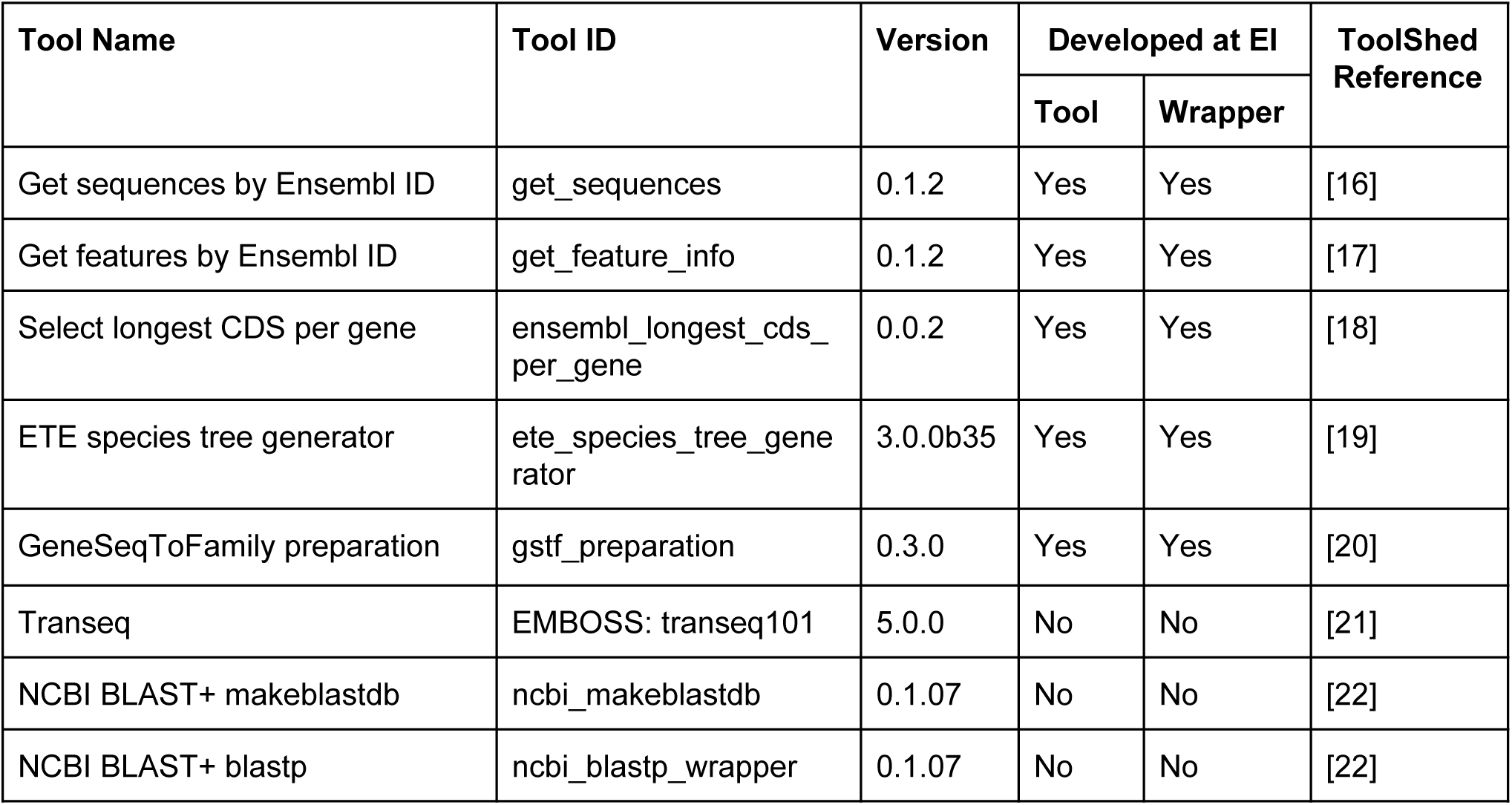
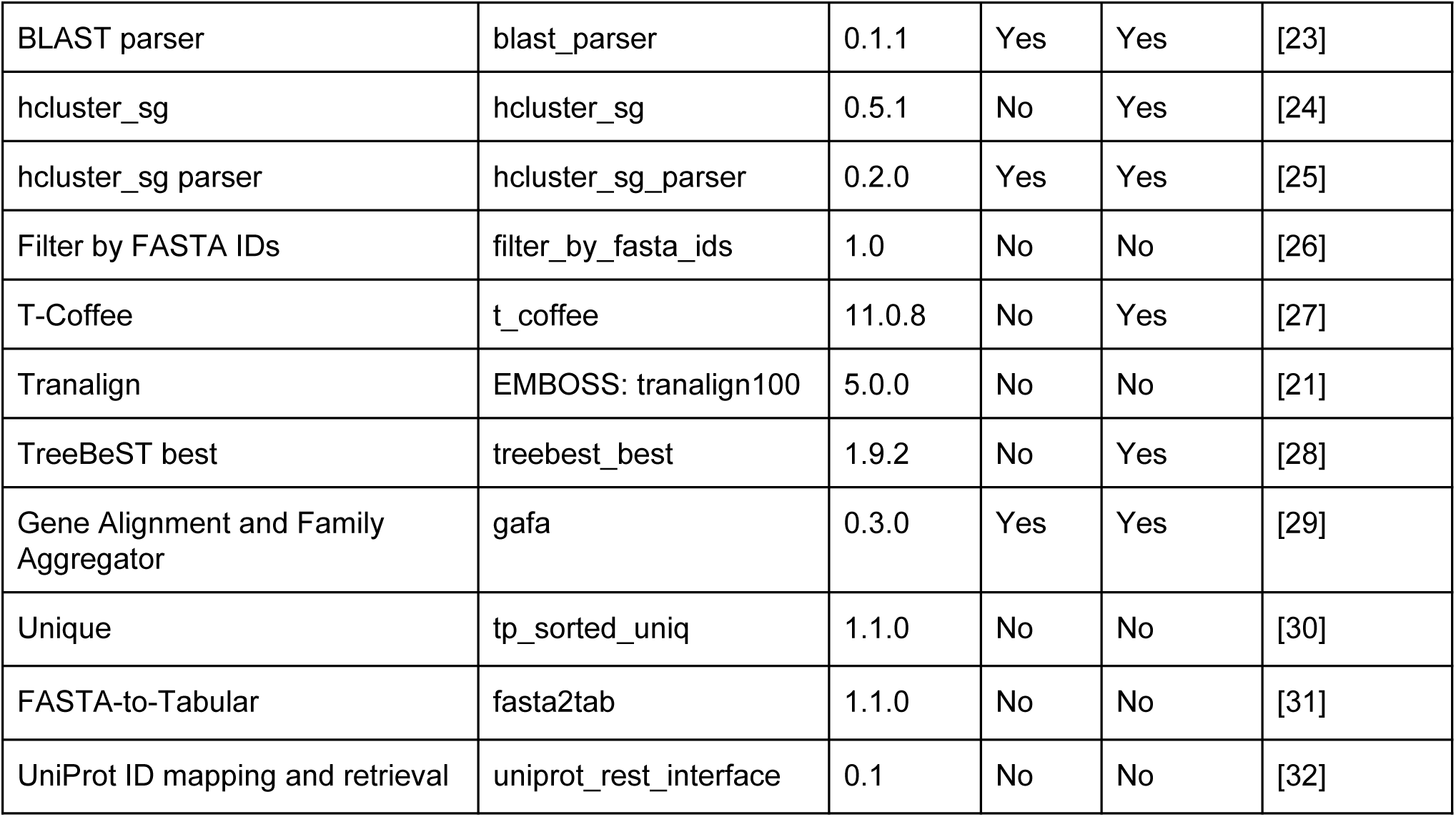
Galaxy tools used in the workflows

The workflow comprises 7 main steps, starting with translation from input coding sequences (CDS) to protein sequences, finding subsequent pairwise alignments of those protein sequences using BLASTP, and then the generation of clusters from the alignments using hcluster_sg. The workflow then splits into two simultaneous paths, whereby in one path it performs the multiple sequence alignment (MSA) for each cluster using T-Coffee, and in the other, generates a gene tree with TreeBeST taking the cluster alignment and a species tree as input. Finally, these paths merge to aggregate the MSA, the gene tree and the gene feature information (transcripts, exons, and so on) into an SQLite [15] database for visualisation and downstream reuse. Each step of the workflow along with the data preparation steps is explained in detail below.

### Data generation and preparation

We have developed a number of tools that assist in preparing the datasets needed by the workflows.

#### Ensembl REST API tools

Galaxy tools were developed which utilise the Ensembl REST API [33] to retrieve sequence information (*Get sequences by Ensembl ID*) and feature information (*Get features by Ensembl ID*) by Ensembl ID from the Ensembl service. REST (REpresentational State Transfer) is an architecture style for designing networked applications [34] which encourages the use of standardised HTTP technology to send and receive data between computers. As such, these tools are designed to help users to retrieve existing data from Ensembl rather than requiring them to manually download datasets to their own computers and then subsequently uploading them into the workflow.

We have also developed:

- *Select longest CDS per gene*, which filters a CDS FASTA file from Ensembl retaining only the longest CDS sequence for each gene
- *ETE species tree generator*, which uses the ETE toolkit [35] to generate a species tree from a list of species names or taxon IDs through NCBI Taxonomy.

### GeneSeqToFamily workflow

#### 0. GeneSeqToFamily preparation

Before GeneSeqToFamily can be run, a data preparation step must be carried out. We have developed a tool called *GeneSeqToFamily preparation* to preprocess the input datasets (gene feature information and CDS) for the GeneSeqToFamily workflow. It converts a set of gene feature information files in GFF3 [36] and/or JavaScript Object Notation (JSON) [37] format to an SQLite database. It also modifies all CDS FASTA header lines by appending the species name to the transcript identifier, as required by *TreeBeST best*. We decided to use an SQLite database to store the gene feature information because:

- the GFF3 format has a relatively inconvenient and unstructured additional information field (9^th^ column)
- searching is much faster and more memory efficient in a database than in a text file like JSON or GFF3, especially when dealing with feature information for multiple large genomes

#### 1. CDS translation

##### Transeq

Transeq, part of the European Molecular Biology Open Software Suite (EMBOSS) [38], is a tool to generate six-frame translation of nucleic acid sequences to their corresponding peptide sequences. Here we use Transeq to convert a CDS to protein sequences in order to run BLASTP and find protein clusters. However, since downstream tools in the pipeline such as TreeBeST require nucleotide sequences to generate a gene tree, the protein sequences cannot be directly used as workflow input and are instead generated with Transeq.

#### 2. Pre-clustering alignment

##### BLAST

This workflow uses the BLAST wrappers [39] developed to run BLAST+ tools within Galaxy. BLASTP is run over the set of sequences against the database of the same input, as is the case with BLAST-all, in order to form clusters of related sequences.

##### BLAST parser

*BLAST parser* is a small Galaxy tool to convert the BLAST output into the input format required by hcluster_sg. It takes the BLAST 12-column output [40] as input and generates a 3-column tabular file, comprising the BLAST query, the hit result, and the edge weight. The weight value is simply calculated as minus log_10_ of the BLAST e-value, replacing this with 100 if this value is greater than 100. It also removes the self-matching BLAST results.

#### 3. Cluster generation

##### hcluster_sg

hcluster_sg performs hierarchical clustering under mean distance for sparse graphs. It reads an input file that describes the similarity between two sequences, and iterates through the process of grouping two nearest nodes at each iteration. hcluster_sg outputs a single list of gene clusters, each comprising a set of sequence IDs present in that cluster. This list needs to be reformatted using the *hcluster_sg parser* tool in order to be suitable for input into T-Coffee and TreeBeST (see below).

##### hcluster_sg parser

*hcluster_sg parser* converts the hcluster_sg output into a collection of lists of IDs, one list for each cluster. Each of these clusters will then be used to generate a gene tree via TreeBeST. The tool can also filter out clusters with a number of elements outside a specified range. The IDs contained in all discarded clusters are collected in separate output dataset. Since TreeBeST requires at least 3 genes to generate a gene tree, we configured the tool to filter out clusters with less than 3 genes.

*Filter by FASTA IDs*, which is available from the Galaxy ToolShed, is used to create separate FASTA files using the sequence IDs listed in each gene cluster.

#### 4. Cluster alignment

##### T-Coffee

T-Coffee is a MSA package, but can also be used to combine the output of other alignment methods (Clustal, MAFFT, Probcons, MUSCLE) into a single alignment. T-Coffee can align both nucleotide and protein sequences [8], and we use it to align the protein sequences in each cluster generated by hcluster_sg.

We modified the Galaxy wrapper for T-Coffee to take a single FASTA (as normal) and an optional list of FASTA IDs to filter. If a list of IDs is provided, the wrapper will pass only those sequences to T-Coffee, which will perform the MSA for that set of sequences, thus removing the need to create thousands of intermediate Galaxy datasets.

#### 5. Gene tree construction

##### Tranalign

Tranalign [38] is a tool that reads a set of nucleotide sequences and a corresponding aligned set of protein sequences and returns a set of aligned nucleotide sequences. Here we use it to generate CDS alignments of gene sequences using the protein alignments produced by T-Coffee.

##### TreeBeST ‘best’

TreeBeST (Tree Building guided by Species Tree) is a tool to generate, manipulate, and display phylogenetic trees and can be used to build gene trees based on a known species tree.

The ‘best’ command of TreeBeST builds 5 different gene trees from a FASTA alignment file using different phylogenetic algorithms, then merges them into a single consensus tree using a species tree as a reference. In GeneSeqToFamily, *TreeBeST best* uses the nucleotide MSAs generated by Tranalign (at least 3 sequences are required) and a user-supplied species tree in Newick format [41] (either produced by a third-party software or through the *ETE species tree generator* data preparation tool, described above) to produce a GeneTree for each family represented also in Newick format. The resulting GeneTree also includes useful annotations specifying phylogenetic information of events responsible for the presence/absence of genes, for example, ‘S’ means speciation event, ‘D’ means duplication, and ‘DCS’ denotes the duplication score.

#### 6. Gene Alignment and Family Aggregation

##### Gene Alignment and Family Aggregator (GAFA)

*GAFA* is a Galaxy tool which generates a single SQLite database containing the gene trees and MSAs, along with gene features, in order to provide a reusable, persistent data store for visualisation of synteny information with Aequatus. *GAFA* requires:

- gene trees in Newick format,
- the protein MSAs in fasta_aln format from *T-Coffee* and
- gene feature information generated with the *GeneSeqToFamily preparation* tool.

Internally, *GAFA* converts each MSA from fasta_aln format to a simple CIGAR string [42]. An example of CIGAR strings for aligned sequences is shown in Figure 3, in which each CIGAR string subset changes according to other sequences.

**Figure 1:**
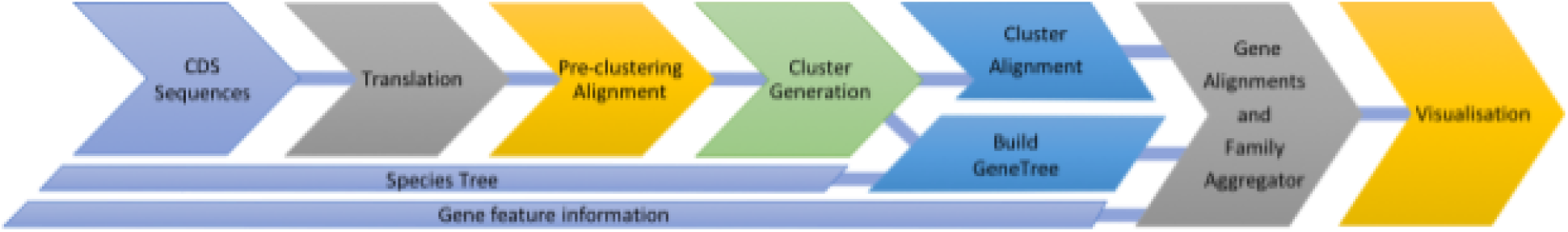
Overview of the GeneSeqToFamily workflow

**Figure 2:**
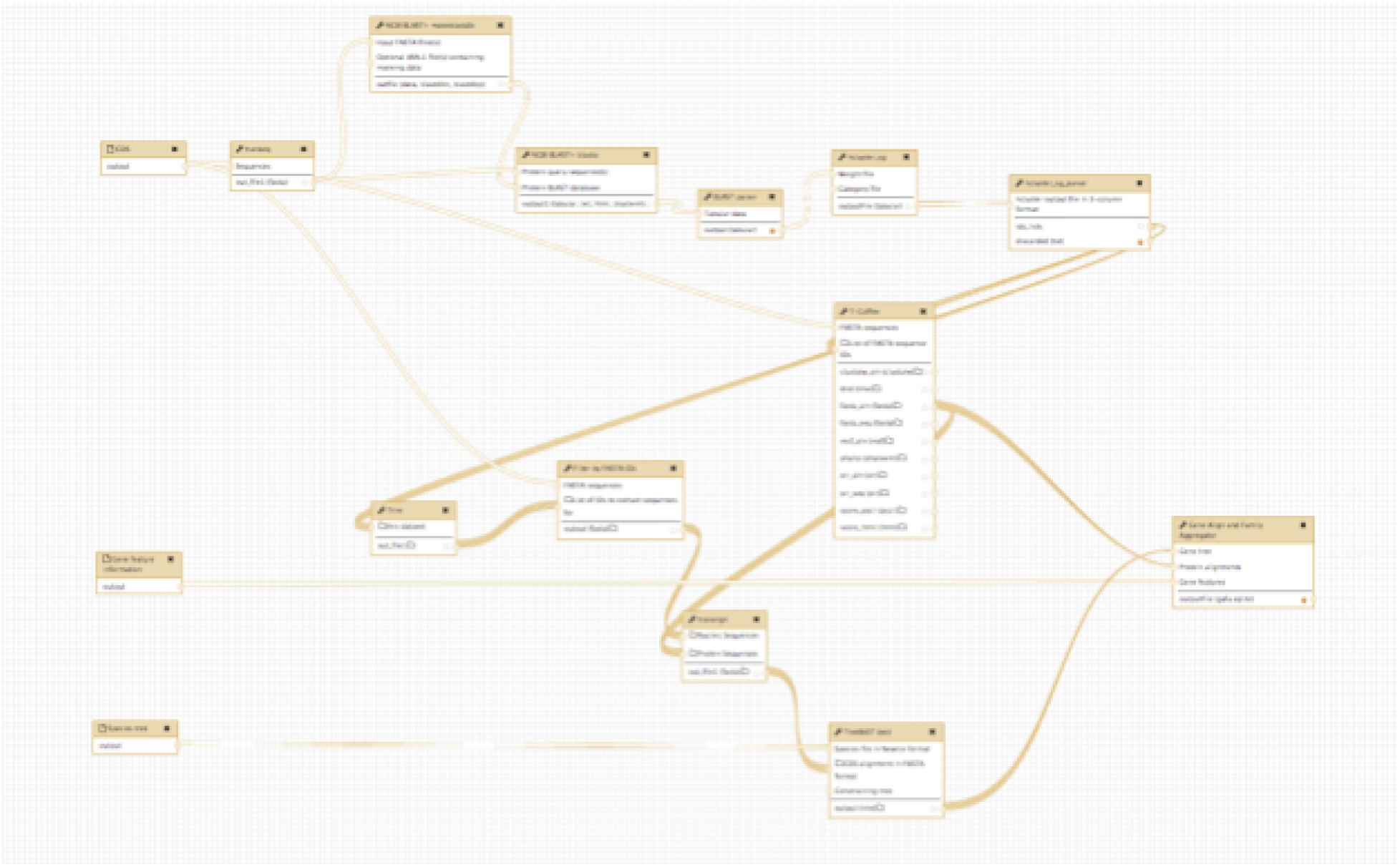
Screenshot from the Galaxy Workflow Editor, showing the GeneSegToFamily workflow

**Figure 3:**
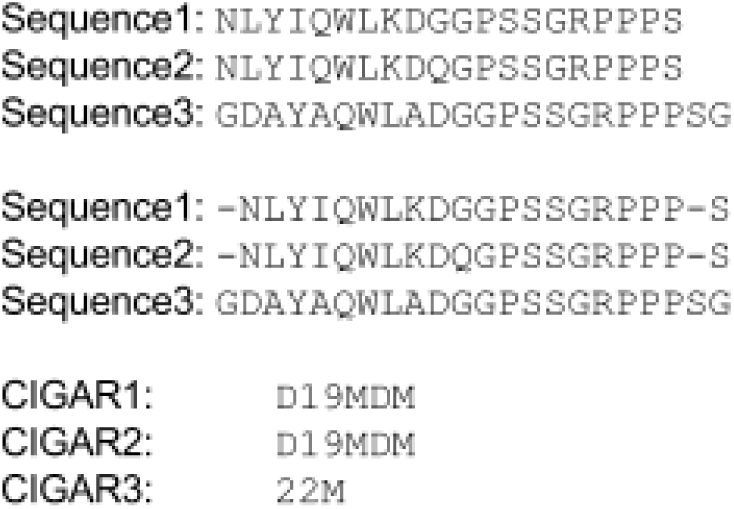
Showing how CIGAR for multiple sequence alignment is generated

The simple schema [43] for the generated SQLite database is shown in Figure 4.

**Figure 4:**
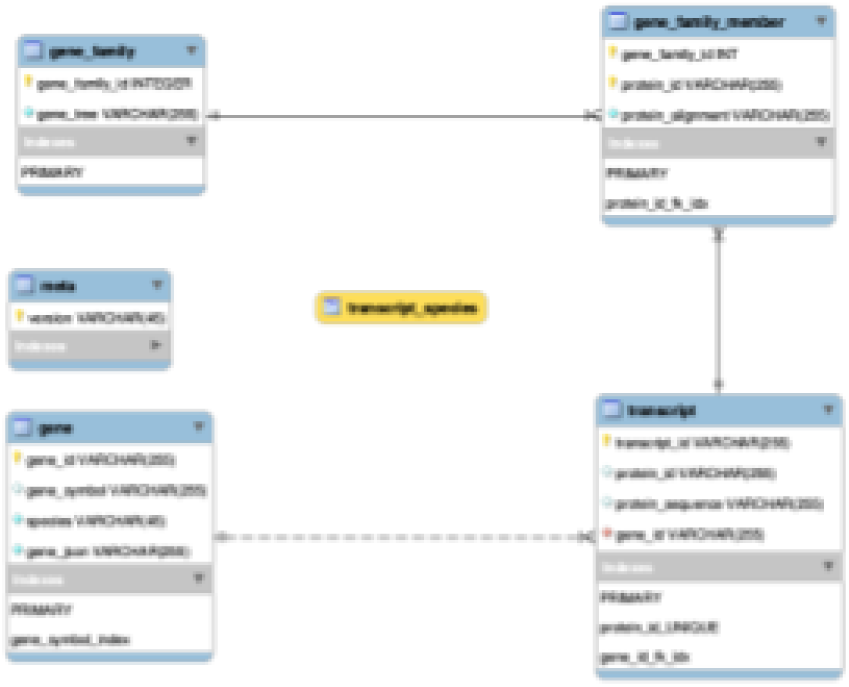
Schema of the GAFA SQLite database

#### 7. Visualisation

##### Aequatus visualisation plugin

The SQLite database generated by the GAFA tool can be rendered using a new visualisation plugin, Aequatus.js. The Aequatus.js library, developed at EI as part of the Aequatus project, has been configured to be used within Galaxy to visualise homologous gene structure and gene family relationships. This allows users to interrogate not only the evolutionary history of the gene family but also the structural variation (exon gain/loss) within genes across the phylogeny. Aequatus.js is available to download from GitHub [43], as visualisation plugins cannot yet be submitted to the Galaxy ToolShed.

### Finding homology information for orphan genes

Although the GeneSeqToFamily workflow will assign most of the genes to orthogroups, many genes within a species might appear to be unique without homologous relationship to any other genes from other species. This observation could be the consequence of the parameters selected, choice of species, incomplete annotations. This could also reflect real absence of homology such as for rapidly evolving gene families. In addition to the GeneSeqToFamily workflow, we also developed two associated workflows to further annotate these genes by:

1. Retrieving a list of orphan genes from the GeneSeqToFamily workflow (see Figure 5) as follows:

a. Find the IDs of the sequences present in the input CDS of the GeneSeqToFamily workflow, but not in the result of *BLAST parser* from the same workflow
b. Add to this list the IDs of the sequences discarded by *hcluster_sg parser*
c. From the input CDS dataset, retrieve the respective sequence for each CDS ID (from the step above) using *Filter by FASTA IDs*

These unique CDS can be fed into the SwissProt workflow below to find homologous genes in other species.
2. Finding homologous genes for some genes of interest using SwissProt (see Figure 6) as follows:

a. Translate CDS into protein sequences using *Transeq*
b. Run BLASTP for the protein sequences against the SwissProt database (from NCBI)
c. Extract UniProt IDs from these BLASTP results, using the preinstalled Galaxy tool *Cut columns from a table* (tool id *Cut1*)
d. Retrieve Ensembl IDs (representing genes and/or transcripts) for each UniProt ID using *UniProt ID mapping and retrieval*
e. Get genomic information for each gene ID and CDS for each transcript ID from the core Ensembl database using *Get features by Ensembl ID* and *Get sequences by Ensembl ID* respectively.

**Figure 5:**
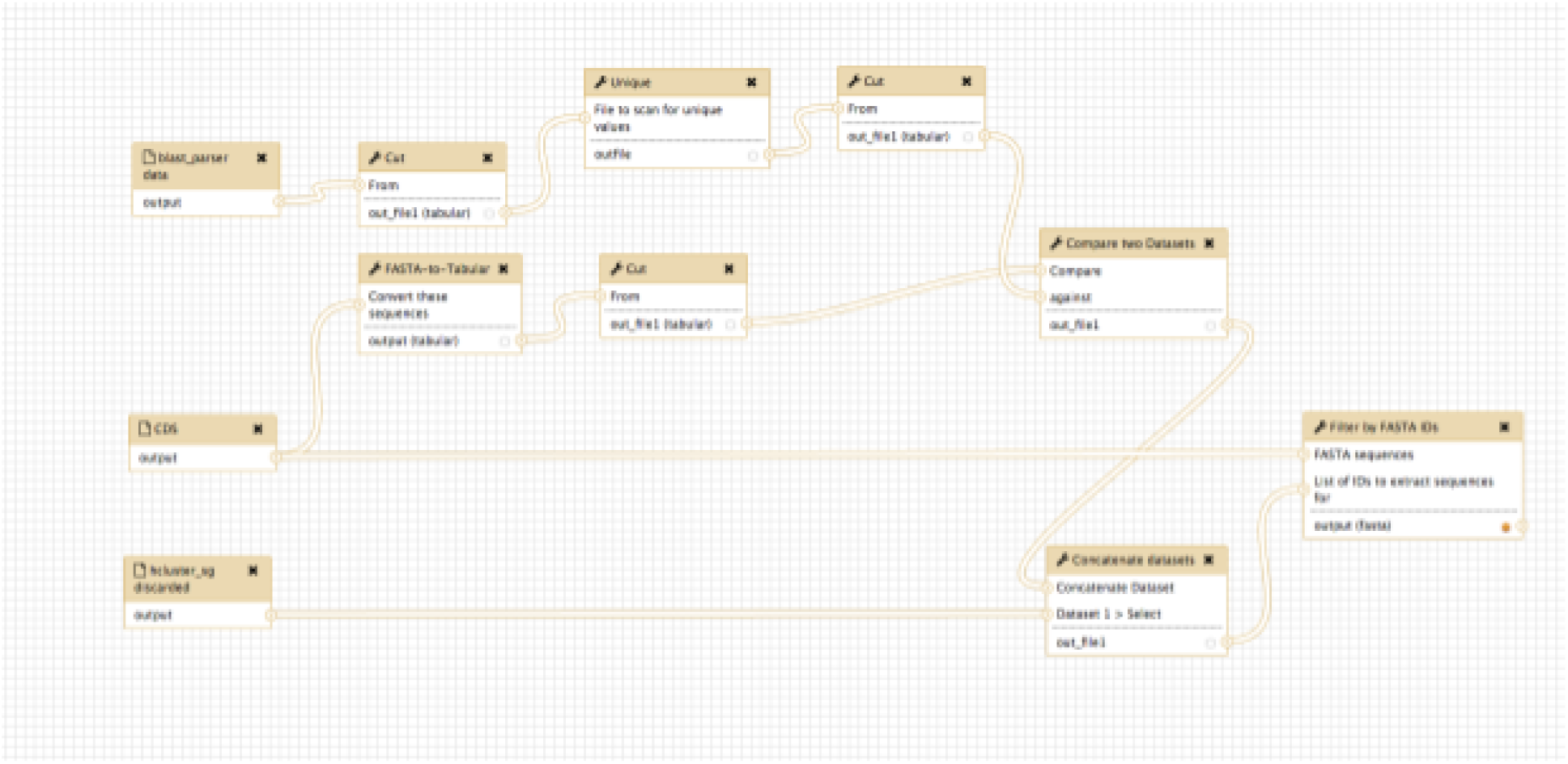
Screenshot from the Galaxy Workflow Editor, showing the orpan gene finding workflow

**Figure 6:**
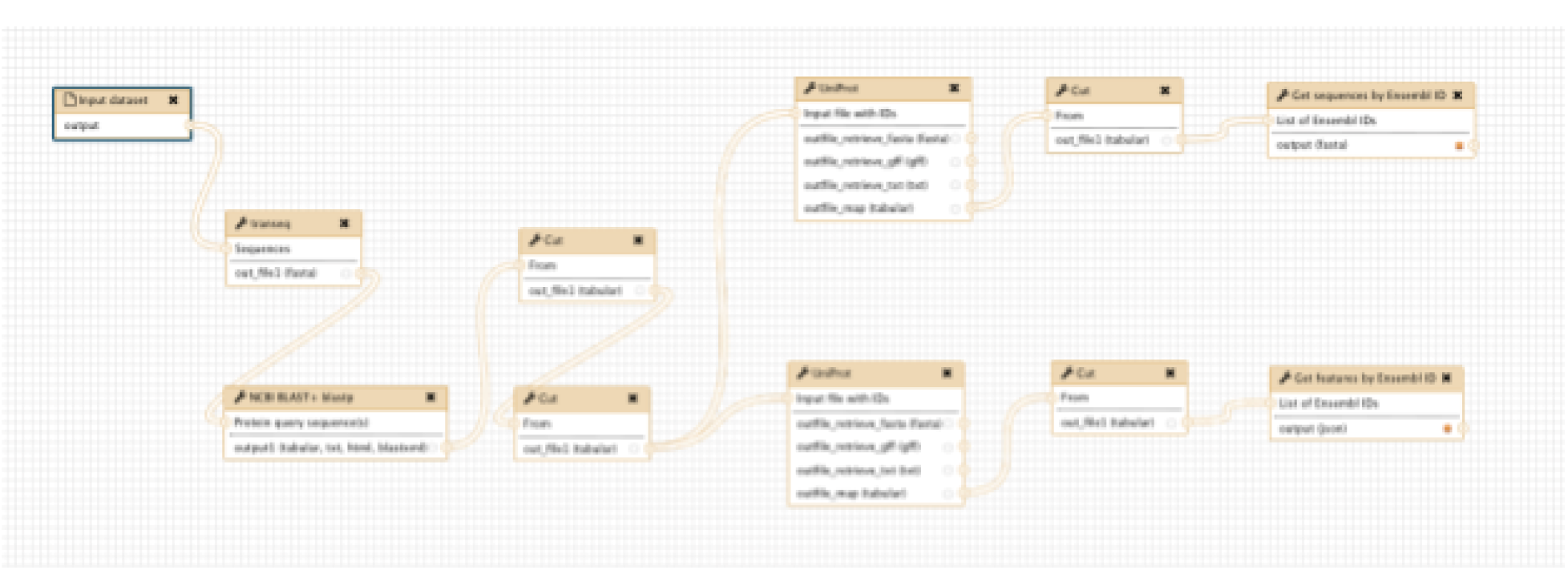
Screenshot from the Galaxy Workflow Editor, showing the SwissProt workflow

The results from this second workflow can be subsequently used as input to GeneSeqToFamily for familial analysis.

## Example use cases

Since BLASTP plays a crucial role in determining gene families, we tested this workflow on a large dataset of CDS from three vertebrate species in order to set a benchmark and find optimum parameters to run the workflow. We downloaded the CDS sequences for *Sarcophilus harrisii* (Tasmanian devil), *Mus musculus* (Mouse), and *Ornithorhynchus anatinus* (Platypus) from Ensembl (release 87) and filtered them to retain only the longest transcript per gene (as in the Ensembl Compara pipeline), obtaining a total of 62,597 CDS. We then ran the GeneSeqToFamily workflow on them using various BLASTP parameters (as shown in Table 3), in order to identify the optimal values for the workflow to generate a gene tree in which members are possibly evolved from a single ancestor gene, and usually with identical biochemical functions such as proteins.

**Table 2:** Results of the GeneSeqToFamily workflow run on 62,597 CDS from 3 species using 6 different BLAST parameter configurations, the complete list of which are shown in Table 3.

| Summary |  |  |  |  |  |  |
| --- | --- | --- | --- | --- | --- | --- |
| Analysis | 1 | 2 | 3 | 4 | 5 | 6 |
| No of families | 8,398 | 13,547 | 13,675 | 12,977 | 13,496 | 17,090 |
| Filtered out (>200) | 12 | 1 | 1 | 1 | 1 | 0 |
| Filtered out (<3) | 2,655 | 2,600 | 3,142 | 1,030 | 2,965 | 5,851 |
| Filtered families to consider | 5,731 | 10,946 | 10,532 | 10,946 | 10,530 | 11,099 |
| Average Family size | 7.15 | 5.11 | 4.50 | 5.11 | 4.50 | 3.65 |
| Median Family Size | 4 | 4 | 3 | 4 | 3 | 3 |
| Largest Family Size | 7,885 | 827 | 599 | 827 | 599 | 64 |
| Smallest Family Size | 1 | 1 | 1 | 1 | 1 | 1 |

**Table 3:** Complete list of BLAST parameter configurations. Analysis 5 is highlighted to denote those parameters that were chosen to be used as workflow defaults.

| Analysis ID | e-value | Max targets per query | Min coverage |
| --- | --- | --- | --- |
| 1 | 1e-03 (Default) | 0 (Default) | 0 (Default) |
| 2 | 1e-03 | 3 | 0 |
| 3 | 1e-03 | 3 | 90% |
| 4 | 1e-10 | 3 | 0 |
| 5 | 1e-10 | 3 | 90% |
| 6 | 1e-10 | 2 | 90% |

Our results show that the number of gene families can vary quite distinctly with different BLASTP parameters. Stringent parameters (Analysis 6) result in a large number of smaller families, while relaxed parameters (Analysis 1) generate a smaller number of large families, which may include distantly related genes. By testing different parameters and comparing the analyses with third party tools such as PantherDB to validate the results against known families, we chose those parameters listed as Analysis 5. These values seem to consistently generate legitimate sets of gene families with closely related family members based on the datasets we tested.

There are caveats, however. BLASTP parameters used in Analysis 5 restrict the maximum number of target sequences per query (max_target_seqs) to 3 (the first hit when using all-versus-all BLAST will always be the query sequence itself). The minimum query coverage per High-scoring Segment Pair (HSP) (qcovhsp) is set to 90% and e-value cut-off to 1e-10, in order to find the HSP closest to the query thus allowing partial matches which could be seen in the event of gene split. If the input CDS contain multiple alternative transcripts per gene, we recommend setting the max_target_seqs parameter to 4 rather than 3 to get a wider range of results from BLAST, thereby helping to generate gene families with matching genes together with alternative transcripts. In contrast setting a value of 3 for the max_target_seqs parameter will restrict the search to only 3 matches per query, and the presence of alternative transcripts will decrease the likelihood of finding matches from other genes, thus increasing the likelihood of splitting of a gene tree into multiple trees and adversely inflating the number of families.

To validate the biological relevance of results from the GeneSeqToFamily workflow, we analysed a smaller set of 23 homologous genes (39 transcripts) from *Pan troglodytes* (chimpanzee), *Homo sapiens* (human), *Rattus norvegicus* (rat), *Mus musculus* (mouse), *Sus scrofa* (pig) and *Canis familiaris* (domesticated dog). These genes are a combination of those found in four gene families, i.e. monoamine oxidases (MAO) A and B, insulin receptor (INSR), BRCA1-associated ATM activator 1 (BRAT1), and were chosen because they are present in all 6 species yet distinct from each other. Though MAO gene variants (A and B) are 70% similar, a single gene tree for all MAO genes could be generated if appropriate parameters are not selected. As such, these genes represent a reliable dataset to test whether the GenSeqToFamily workflow can reproduce already known gene families.

Before running the workflow, feature information and CDS for the selected genes were retrieved from the core Ensembl database using the helper tools described above (*Get features by Ensembl ID* and *Get sequences by Ensembl ID* respectively). A species tree was generated using *ETE species tree generator* and CDS were prepared with *GeneSeqToFamily preparation*. We ran the GeneSeqToFamily workflow on these data using the parameters shown in Analysis 5 of Table 3, but we set max_target_seqs as 4 (as described in the previous use case) to get a wider range of results from BLAST because our dataset includes alternative transcripts. This workflow generated 4 different gene trees, one for each gene family. Figure 7, 8, 9 and 10 show the resulting gene trees for MAOA, MAOB, BRAT1 and INSR gene families. Different colours of the nodes in each gene tree on the left-hand-side highlight potential evolutionary events, such as speciation, duplication, and gene splits. Homologous genes showing shared exons use the same colour in each representation, including insertions (black blocks) and deletions (red lines). The GeneTrees for these genes are already available in Ensembl and we used them to validate our findings [44] [45] [46] [47]. Our gene trees exactly matched the Ensembl GeneTrees, showing that the workflow generates biologically valid results. We have provided the underlying data for this example along with the submitted workflow in figshare [48].

**Figure 7:**
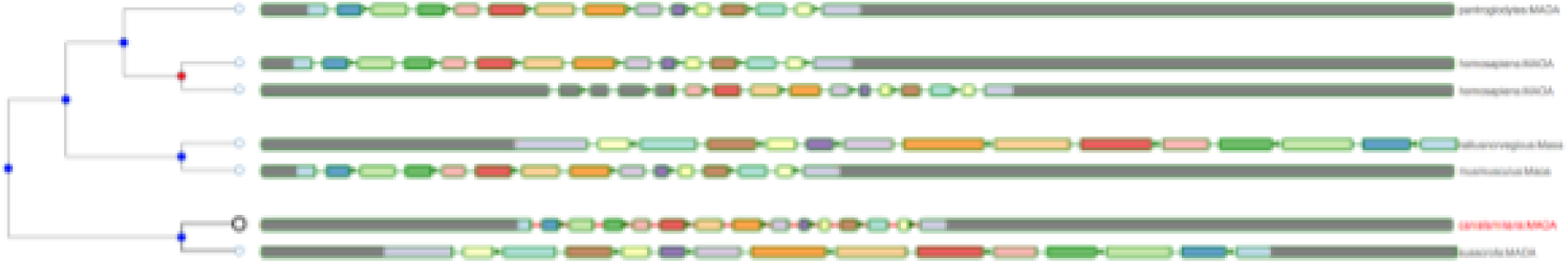
Homologous genes of MAOA of *Canis familiaris from Mus musculus, Pan troglodytes, Homo sapiens. Rattus norvegicus, Sus scrofa and Canis familiaris.*

**Figure 8:**
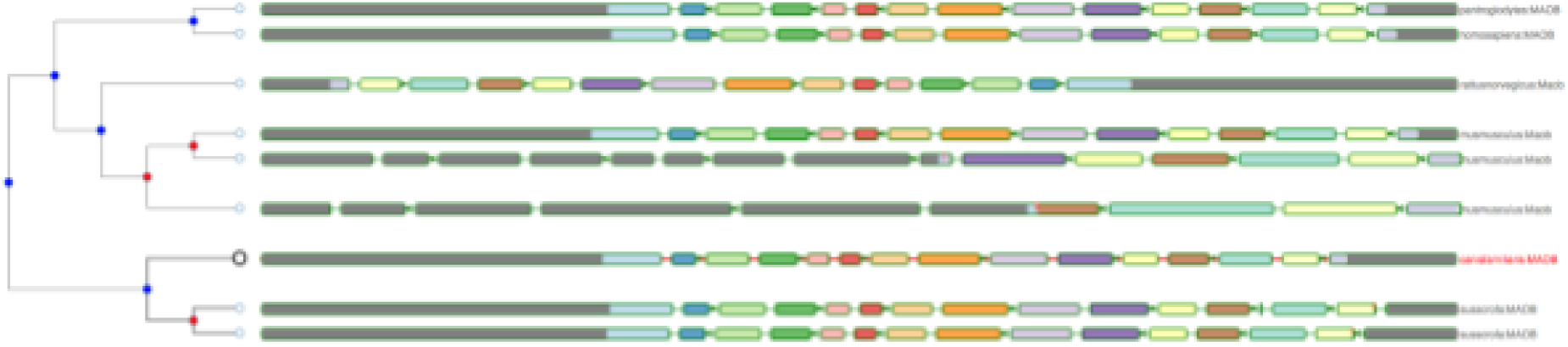
Homologous genes of MAOB of *Canis familiaris from Mus musculus, Pan troglodytes, Homo sapiens. Rattus norvegicus, Sus scrofa and Canis familiaris.*

**Figure 9:**
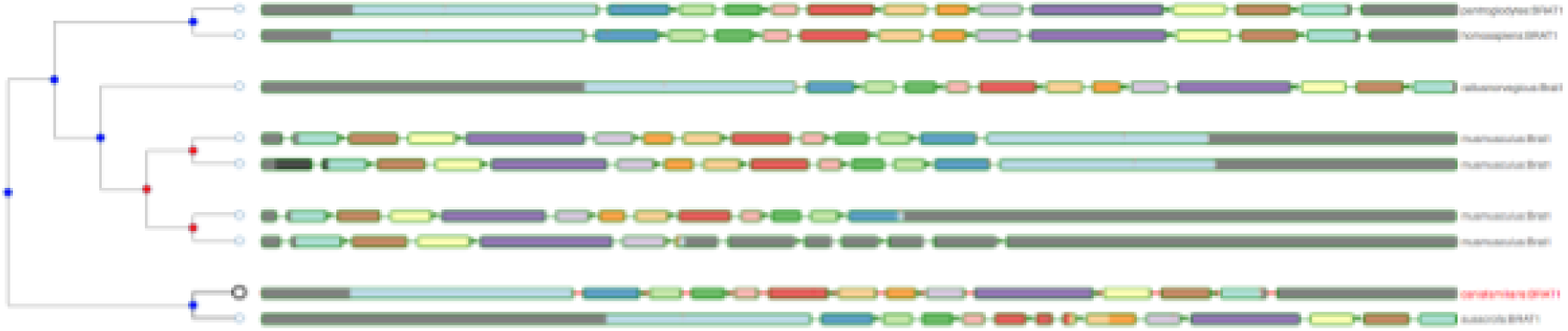
Homologous genes of BRAT1 of *Canis familiaris from Mus musculus, Pan troglodytes, Homo sapiens. Rattus norvegicus, Sus scrofa and Canis familiaris.*

**Figure 10:**
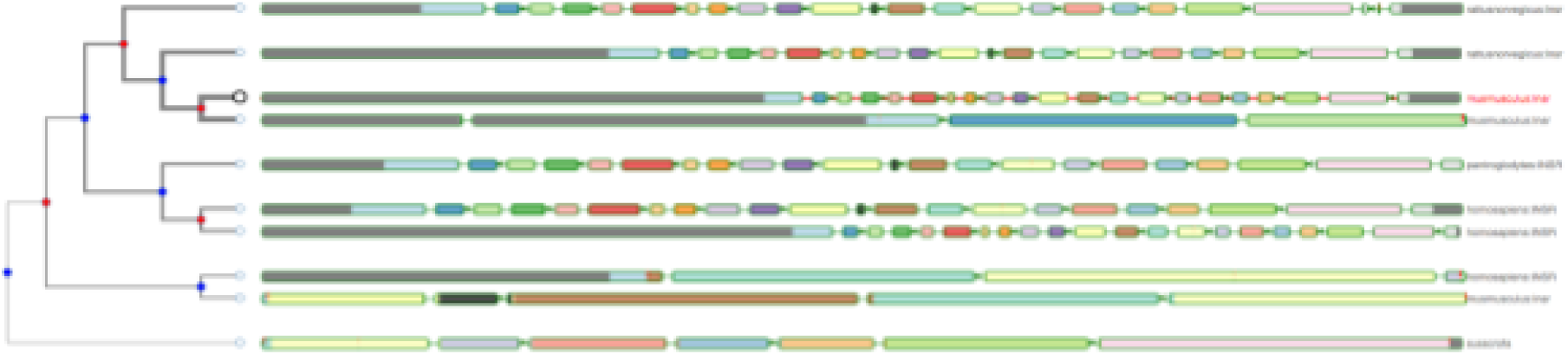
Homologous genes of INSR of *Canis familiaris from Mus musculus, Pan troglodytes, Homo sapiens. Rattus norvegicus, Sus scrofa and Canis familiaris.*

## Conclusion

The ultimate goal of the GeneSeqToFamily is to provide a user-friendly workflow to analyse and discover homologous genes using the Ensembl Compara GeneTrees pipeline within the Galaxy framework, where users can interrogate genes of interest without using the command-line whilst still providing the flexibility to tailor analysis by changing configurations and tools if necessary. We have shown it to be an accurate, robust, and reusable method to elucidate and analyse potentially large numbers of gene families in a range of model and non-model organisms. The workflow stores the resulting gene families into a SQLite database, which can be visualised using the Aequatus.js interactive tool, as well as shared as a complete reproducible container for potentially large gene family datasets.

Gradually, we hope that the Galaxy community will undertake their own analyses and feedback improvements to various tools, and publish successful combinations of parameters used in the GeneSeqToFamily workflow. We encourage this process by allowing users to share their own version of GeneSeqToFamily workflow for appraisal by the community.

### Future directions

In terms of core workflow functionality, we would like to incorporate pairwise alignment between pairs of genes for closely related species in addition of the MSA for the gene family, which will help users to compare orthologs and paralogs in greater detail.

We also plan to include explicit integration of the PantherDB resources [49]. Protein ANalysis Through Evolutionary Relationships (PANTHER) is a classification system to characterise known proteins and genes in order to certify genomic annotation. Association of PantherDB with GeneSeqToFamily will enable the automation of gene family validation and add supplementary information about those gene families, which could then be used in turn to further validate novel genomics annotation.

We also plan to add the ability to query the *GAFA* SQLite database using keywords, to make it easy for users to find gene trees which include their genes of interest without needing to delve into the database itself.

## Availability and requirements

**Project name:** GeneSeqToFamily

**Project home page:** https://github.com/TGAC/earlham-galaxytools/tree/master/workflows/GeneSeqToFamily

**Archived version:** 0.1.0

**Operating system(s):** Platform independent

**Programming language:** JavaScript, Perl, Python, XML, SQL

**Other Requirements:** Web Browser; for development: Galaxy

**Any restrictions to use by non-academics:** None

**License:** The MIT License

## Availability of supporting data

The example files and additional data sets supporting the results of this article are available in figshare [48].

## Acknowledgements

AT, WH and RPD are supported by BBSRC institute strategic programme grant funds awarded to EI. NS is funded under the BBSRC Biomathematics and Bioinformatics Training fund (2014). This research was supported in part by the NBI Computing infrastructure for Science (CiS) group who provide technical support and maintenance to EI’s High Performance Computing cluster and storage systems, enabling us to develop this workflow.

We would like to thank Matthieu Muffato from the European Bioinformatics Institute (EBI) for his advices during the initial stage of the project.

